# Branch-recombinant Gaussian processes for analysis of perturbations in biological time series

**DOI:** 10.1101/291757

**Authors:** Christopher A. Penfold, Anastasiya Sybirna, John Reid, Yun Huang, Lorenz Wernisch, Zoubin Ghahramani, Murray Grant, M. Azim Surani

## Abstract

**Motivation:** A common class of behaviour encountered in the biological sciences involves branching and recombination. During branching, a statistical process bifurcates resulting in two or more potentially correlated processes that may undergo further branching; the contrary is true during recombination, where two or more statistical processes converge into one. A key objective is to identify the time of this bifurcation (branch time) from time series measurements e.g., comparing a control time series with a perturbed time series. Whilst statistical treatments for the two branch (control versus treatment) case exists, the ability to infer more complex branching structure from time series data remains open. Gaussian processes (GPs) represents an ideal framework for such analysis, allowing for nonlinear regression that includes a rigorous treatment of uncertainty. Currently, however, GP models only exist for two-branch systems. Here we highlight how arbitrarily complex branching processes can be built using the correct composition of covariance functions within a GP framework, thus outlining a general framework for the treatment of branching and recombination in the form of branch-recombinant Gaussian processes (B-RGPs). We first demonstrate the performance of B-RGPs compared to a variety of existing regression approaches, and demonstrate robustness to model misspecification. B-RGPs are then used to investigate the branching patterns of *Arabidopsis thaliana* gene expression following inoculation with the hemibotrophic bacteria, *Pseudomonas syringae DC3000*, and a disarmed mutant strain, hrpA. By grouping genes according to the number of branches, we could naturally separate out genes involved in basal immune response from those subverted by the virulent strain, and show enrichment for targets of pathogen protein effectors. Finally, we identify two early branching genes WRKY11 and WRKY17, and showed that groups of genes that branched at similar times to WRKY11/17 were enriched for W-box binding motifs, and overrepresented for genes differentially expressed in WRKY11/17 knockouts, suggesting that branch time could be used for identifying direct and indirect binding targets of key transcription factors. **Software is available from:** https://github.com/cap76/BranchingGPs.

## 1 Introduction

A common class of behaviour encountered in the biological sciences involves branching. In a branching process, often driven by a biological perturbation, a statistical process bifurcates at a specific time, leading to two potentially correlated processes that may, themselves, undergo further branching (Poincaré 1885). Reciprocal behaviour is encountered in recombination processes, where two or more statistical processes converge.

Such branching and recombination is frequently encountered in transcriptional time series data involving host-pathogen interactions. The initial response to infection is the activation of innate immunity, a highly-conserved response based upon perception of non-self. Subsequently, pathogens can deliver protein effectors which collectively suppress immunity, and later collaborate to reconfigure plant metabolism for pathogen nutrition. Thus, initially, the expression dynamics of key infection marker genes will be identically distributed in both infected and uninfected host cells. Expression patterns will begin to diverge as the host mounts immunity; in many cases, this innate immune response is suppressed by the pathogen, potentially driving expression levels of certain genes back to uninfected levels. Indeed nearly 50% of the transcriptome is observed to be differentially expressed during some plant infections (Windram et al. 2012; Lewis et al. 2015). More complex patterns of branching and recombination may exist in such datasets due to the ongoing evolutionary arms race between pathogens and their hosts (Jones and Dangl 2006; Boller and He 2009).

The ability to infer the timing of bifurcations in individual genes should reveal important information about the onset and development of infection. The inference of branching and recombination processes from systems level measurements, such as collections of microarray or RNA-sequencing data, remains a difficult challenge, partially due to datasets being noisy in nature, with (potentially) missing observations or uneven temporal sampling. The dynamic nature of different biological systems may also vary significantly, frustrating efforts to find a robust, broadly applicable approach to the inference of branching and recombination. Nonparametric Bayesian approaches to inference would therefore be advantageous, addressing these key issues. Gaussian processes represent a flexible nonparametric Bayesian approach to nonlinear regression able to gracefully cope with uncertainty, uneven sampling and a diverse range of dynamic behaviour (Rasmussen and Williams 2006). However, currently, Gaussian processes treatments for branching processes have only been developed for the two-branch case (Yang et a1. 2016; Stegle et al. 2010). Here we develop an approach to inference for arbitrarily complex branching and recombination processes, in the form of branch-recombinant Gaussian processes (B-RGPs). In Section 2 we first introduce B-RGPs, highlighting their key limiting behaviour. In Section 3 we demonstrate the advantages of B-RGPs over GPs on a variety of simulated datasets, and in Section 4 we demonstrate the utility of our approach on genome-scale time-course microarray data by identifying transcriptional branching and recombination in *Arabidopsis thaliana* infected with bacterial pathogen *Pseudomonas syringae.* Finally, in Section 5, we discuss a variety of possible applications for B-RGPs and future avenues for research.

### 2 Methods

Within a Bayesian setting, Gaussian processes (GPs) can be used to represent prior distributions over smooth functions, providing a flexible framework for regression and classification with robust treatment of uncertainty (Rasmussen and Williams 2006). This makes GP-based approaches ideal frameworks for quantifying the dynamics of gene expression from biological observations (Stegle et al. 2010; Kalaitzis and Lawrence 2011; Breeze et al. 2011; Hensman, Lawrence, and Rattray 2013). For regression, we typically have a set of observations, ***y***, assumed to be noisy instances of a continuous underlying function at input locations ***t***:***y** = f*(***t***) + *∈*, where ε represents Gaussian additive noise. In our applications, ***y*** will typically be used to denote a vector of the observed expression levels for a given gene at times, ***t***. We can assign the unknown function a GP prior, denoted *f*(*t*) ~ 𝒢𝒫(μ(*t*))*k*(*t,t'*)), and analytically evaluate the posterior distribution at a set of new input locations, ***t*\***. The marginal likelihood, too, may be analytically evaluated, making GPs a flexible and efficient framework for both prediction and model comparison. Previous GP-based approaches to branching have been outlined for the two-dataset case i.e., where there exists two biological processes following branching. These include the studies by Stegle et al. (2010), who developed a GP two-sample approach, based on mixtures of GPs, and the more recent work of Yang et al. (2016), who demonstrate explicitly how a two-branch process can be encoded within a joint GP model. To our knowledge, the generalisation of GPs to more than two branches has not been addressed, whilst no explicit closed-form solution to recombination has been outlined.

A useful extension to the GP framework is the multiple output hierarchical Gaussian process (HGP; Hensman, Lawrence, and Rattray (2013)), in which a basal process is defined by a zero-mean GP with covariance function *k*_1_(*t,t'*), with a subsequent process having mean *f*_1_(*t*) and covariance function, *k*_2_(*t,t'*):

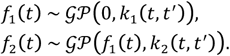

Within this framework, we assume noisy observations of the functions, *y*_1_=*f*_1_(*t*) + *∈*, and *y*_2_ *=f*_2_(*t*) *+∈* and may analytically evaluate the posterior distribution at a new set of input locations for prediction, or the marginal likelihood for model comparison. A class of branching behaviour can naturally be encoded within this HGP framework, assuming the basal (main branch) process is defined by a zero-mean GP with covariance function *k*_*b*_1__ (*t,t'*), with a subsequent process having mean *f*_*b*_1__(*t*)and an appropriate covariance function that ensures the two processes are identically distributed prior to an arbitrarily chosen time point, *t*_*b*_:

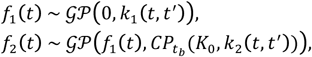

where *K*_0_= *K*_0_(*t,t'*) denotes a zero-kernel, and 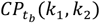 denotes a change-point kernel (Lloyd et al. 2014), defined as:

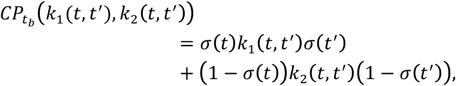

where 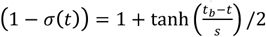. Here we introduce two hyperparameters: *t*_*b*_, which represents the branch time, and *s*, which controls how fast data belonging to the second branch transition from the basal process to the branch process. Note that each data point must be assigned a branch label, *z* ∈ [1,2], according to which branch it belongs to e.g., *z* = 1 will be used to denote data belonging to the control or wildtype branch, with z = 2 referring to the perturbed dataset. For a two branch case observations are *a priori* Gaussian distributed, ***y***_1_,***y***_2_|***t,z**~ N*(**0**,*K*(***t, t'***,*z,z'*)), where:

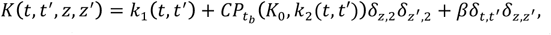

and the delta function *δ*_*z,*2_*δ*_*z′,*2_ ensures the change-point kernel only operates over the second branch i.e., where the branch label *z* = 2 and *z′* = 2. Within this framework, we may again make a prediction ***y\**** at a new input location, **(*t*,z\**),** and analytically evaluate the marginal likelihood, allowing us to compare the goodness of fit between different branching processes.

We can allow further branches that independently diverge from the main branch, with each data point assigned a branch label. For a n-component system *z* ∈ [1,…, *n*] and we have the following covariance function:

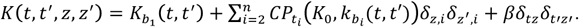

Alternatively, rather than each branch diverging from the main process, each branch could itself give rise to further branches in a recurrent manner e.g., a basal (main branch) from which a secondary branch diverges, with a third branching from the second and so forth. For an n-component recurrent branching system we have:

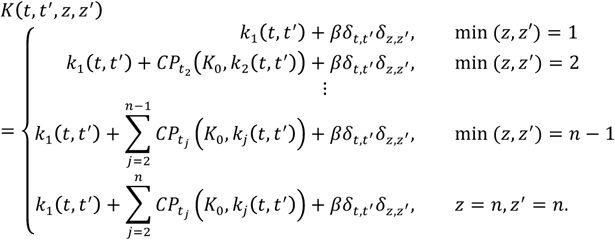

When observation data for all branches are specified over identical time points, the covariance matrix can be expressed in a more compact notation:

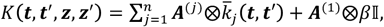

where ⊗ denotes the Kronecker product and:

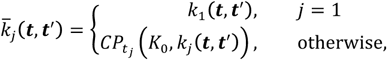

where *m* represents the number of unique time points, 𝕀 represents an (m×m) identity matrix, and ***A***^(*j*)^ = ***u u***^T^, with ***u*** representing a column vector of length *n*, with ones in elements *j* through *n* and zeros every-where else. Far more complex branching patterns can easily be built via the correct composition of independent and recurrent branching covariance functions.

As well as building branching structures of arbitrary complexity, we further note that the dynamic behaviour of the individual branches themselves may themselves be arbitrarily complex, comprised of any linear combination of positive semi-definite kernels. In Supplementary Figure 1(a) we indicate example behaviour of simple branching GPs.

**Figure 1:**
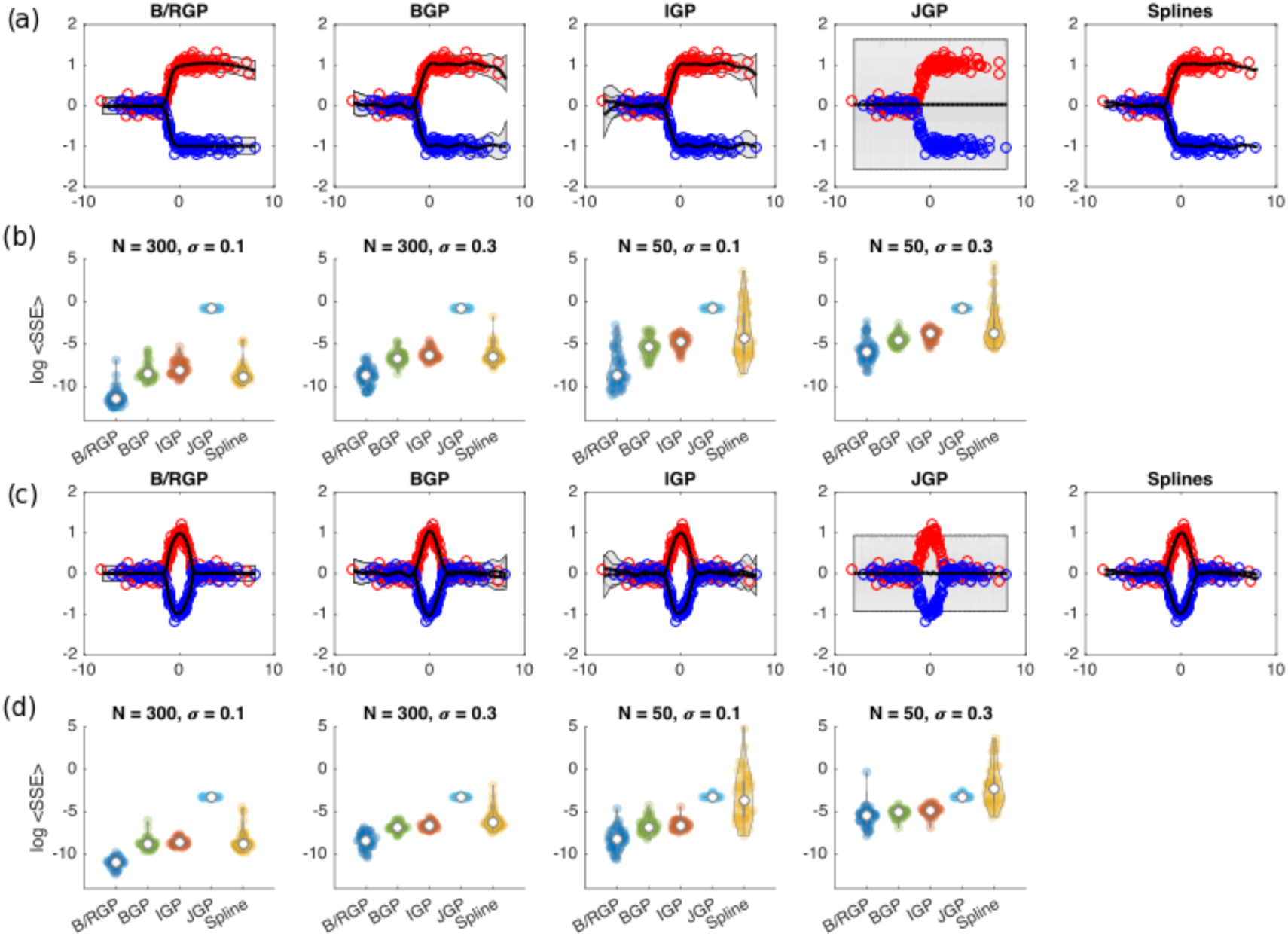
(a) Fits to a two-component branching process using a branch GP outlined here, the branching GP outlined in Yang et al. (2016), independent GPs, a joint GP and independent splines. (b) We indicate the log mean sum squared error for each of the methods for different number of training points and for different noise levels. (c) Fits to a two component branch-recombinant process using branch-recombinant GPs, branch GPs of Yang et al. (2016), independent GPs, joint GPs and splines. (d) Log mean sum squared error for the different approaches for different number of data points and noise levels.

### 2.1 Recombinant Gaussian processes

Recombinant processes can be defined in a reciprocal fashion to branching processes. Notable examples might include the reprogramming of different terminally differentiated cell lineages to iPSCs (Gurdon 1962; Takahashi and Yamanaka 2006). We can describe a two-component system via the following composition of covariance functions:

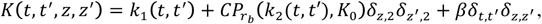

which encodes the main branch process, with a second (potentially correlated) process that recombines after time *r*_*b*_. Multiple processes can again be allowed to independently recombine with the main branch, or recurrently recombine via a series of parental branches, analogously to branching GPs. Example recombinant GPs are shown in Supplementary Figure 1(b).

### 2.2 Branch-recombinant Gaussian processes

Another important process exists where a statistical process transiently branches into two or more processes, before recombining back into a single process. Such combinations of branching and recombination may be encountered during development when where there exists more than one route to a terminal cell fate, as may be the case in certain neuronal lineages (Zawadzka et al. 2010), as well as in certain diseases, such as during dedifferentiation of cancer cells (Friedmann-Morvinski and Verma 2014). An example two-component system can be encoded by the following covariance function:

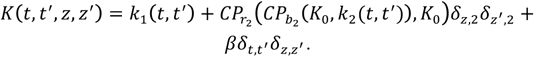

Again, more complex patterns, with arbitrary numbers of branches and recombination, can readily be built with GPs via the correct composition of covariance functions, with more complex examples shown in Supplementary Figure 1(c).

### 2.3 Optimisation, run time and limiting behaviour

A key advantage of the B-RGP framework outlined here over existing approaches (Yang et al., 2016) is the ability to fit arbitrarily complex branch-recombinant structures i.e., more branches. Unlike the earlier work of Yang *et al.* (2016), all hyperparameters including those relating to branch and recombination times can be directly optimised via gradient based approaches e.g., type II ML estimators. In general we note that inference with B-RGPs scales as any other GP, with complexity *𝒪*(*n*^3^), where *n* is the number of observations, although sparse approximations are possible. The time required for optimisation of hyperparameters via type II ML estimates varied: for a dataset with 300 observations, 1000 steps of the gpml minimize function took approximately 30 seconds on a Desktop computer (2.5 GHz Intel Core i7), although it should be noted that, in many cases, full convergence could require more than 1000 steps. This makes B-RGPs slightly slower than the time taken for DEtime (Yang et al., 2016), which, for the same dataset and default parameters, ran in around 10 seconds.

Depending upon the branch time hyperparameters and other hyperparameters in the change-point kernel, the behaviour of B-RGPs can naturally tend towards either an independent GP or a HGP. Specifically, for a branching GP, when *σ*(*t*)→ 1, as may be the case when (*t_b_ – t*)*/s* is very large, such as when a branch occurs much later than the last data point, then a BGP will behave as single joint GP with behaviour defined by the main branch kernel only. When *σ*(*t*)→ 0, as may be the case when (*t_b_ – t*)*/s* has increasingly low values, such as when branching occurs much earlier than the first observations, then the BGP will behave as a HGP. Similar limiting behaviour applies for recombination processes.

### 3 Results

As a preliminary test of the B-RGP framework we fitted five simulated labelled time series datasets, and evaluated the predictive accuracy over a range of test locations, comparing the accuracy to that achieved using DEtime (Yang *et al.*, 2016), independent Gaussian process regression (IGP) over the individual branches, joint Gaussian process regression (JGP) over the union of data, and splines. We first evaluated the ability to fit the following branching process:

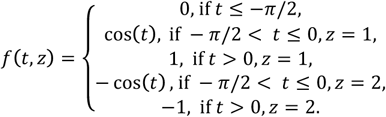

where z indicates the branch label. Random input locations were sampled, **t**~𝒩(**0**, 3𝕀), with branch labels assigned with equal probability, *z_i_*∈ [1,2]. Observations were generated as noisy instances, 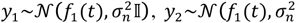, where *σ_n_*∈ [0.1,0.3]. A three-component branching process, comprised of a (latent) main process from which two observed branches diverge, was fitted to the simulated data, with hyperparameters optimised using type II maximum likelihood (ML). The base kernel and all kernels were set a squared-exponentials. Branch time hyperparameters were tied i.e., *t_b_*_1_ *= t_b_*_2_, with initial values set as *t_b_*_1_= 4, log *S_b_*_1_,*b*_2_= 0.5, *σ*_*n*_=0.2, and all other hyperparameters *θ* =[*l*_*b*_0__,*σ*_*b*_0__,*l*_*b*_1__,*σb*_1_,*lb*_2_,*δ*_*b*_2__] initiated as i.i.d. random variables *θ_i_~U*(0.1,1). In Figure 1(a) we indicate an example posterior fits to the data using a BGP, IGPs, JGPs, and splines respectively. In Figure 1(b) we indicate the log mean sum squared error (SSE) over 50 randomly initiated runs using N=50 and N=300 training points and for different noise levels. The B-RGPs shows superior fits (reduced SSE) and decreased negative log marginal likelihood compared to other approaches. The fits obtained using DEtime also appeared to perform well in all cases, outperforming independent GPs, and demonstrating the usefulness of using more accurate generative models for inference of branching data. Next we evaluated the ability of branching GPs to estimate the branch time. In Supplementary Figure 2(b) we plot the branch time versus inferred branch time for 50 instances and compare to that achieved using DEtime (Yang *et al.*, 2016). We note that the correlation for our approach (R=0.9999) indicates good ability to infer branch times, and was greater than that the correlation when using DEtime with default settings (R=9007). Here the increased accuracy partly comes from the ability to directly optimise the branch time hyperparameters via type II ML estimates, rather than relying on a grid search of inferred branch times. To further explore the ability to infer branch times for datasets with missing observations, we repeated this experiment, but excluded observations close to the true branch point, specifically removing any data points where 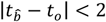, where, 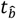 represents the true branch time, and *t*_0_ is the time of the data point. Even with missing observations centred at the true branch time, the inferred branch times were found to be highly correlated with the true branch time (R=0.9649; Supplementary Figure 2(c)).

**Figure 2:**
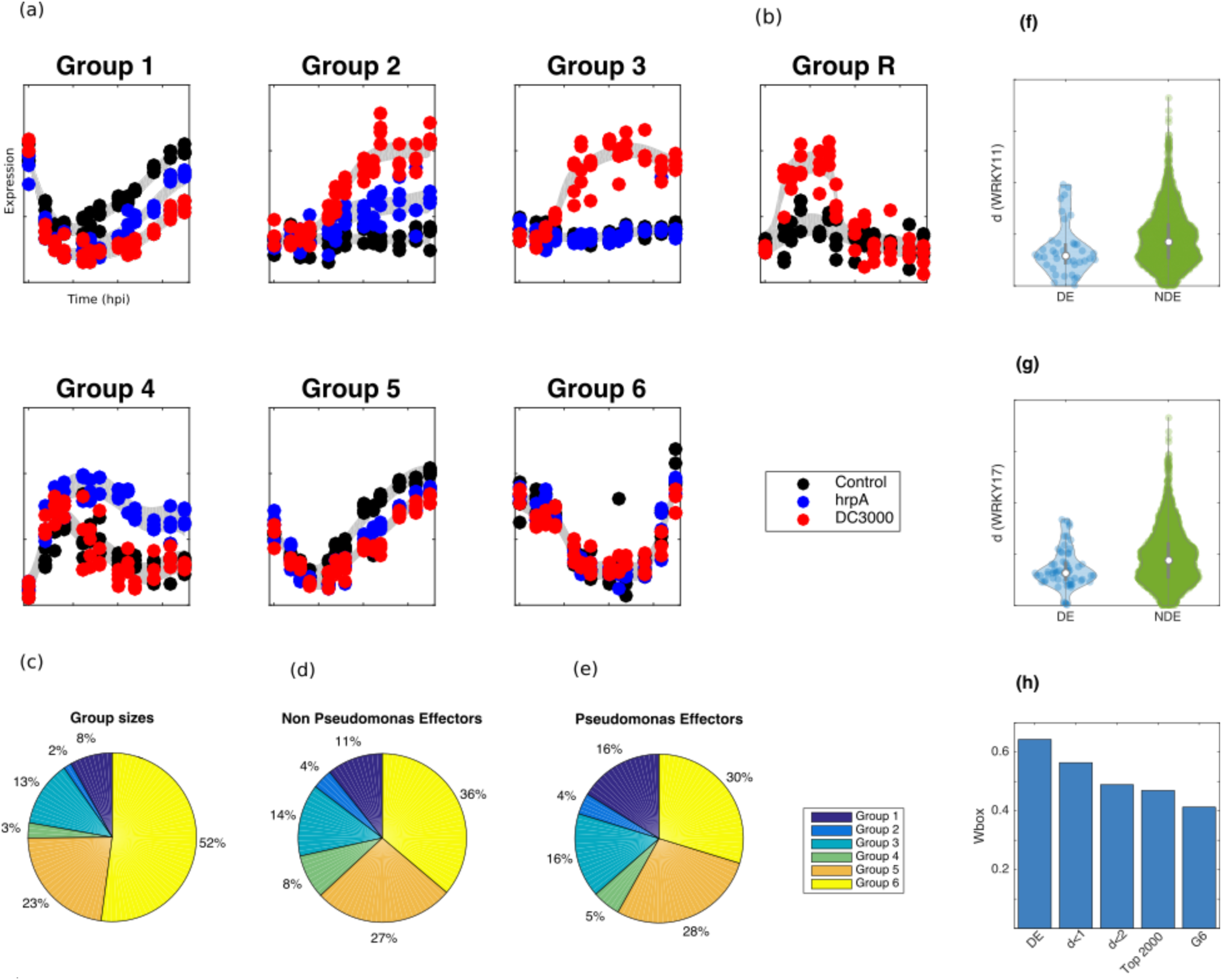
Branching processes were fitted to the three *Arabidopsis* time series, with hyperparameters optimised to MAP values, and the BIC used to select optimal branching structure. **(a)** Example expression profile plots for each of the different classes of branching. **(b)** Example expression profile of a branch-recombinant structure within the dataset. **(c)** The prevalence of each of the six groups within the dataset, compared to the breakdown of *non-Pseudomonas* effector targets **(d)**, and *Pseudomonas-*effector targets show a clear enrichment of effector genes **(e)**. **(f, g)** The Euclidean distance of branching times of genes from that of WRKY11/17 is statistically lower in genes that are DE in WRKY11/17 knockouts versus those that are NDE, indicating that perturbation times are predictive of direct and indirect targets of WRKY11/17. **(h)** The prevalence of Wbox motifs decreases amongst sets of genes whose branch times are increasingly distant from WRKY 11.

In dataset 2, we assumed the following branch-recombinant process:

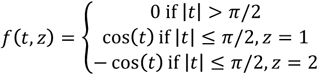

where z indicates the branch label. Again, randomly determined input locations were sampled as before. A three-component branch-recombinant GP comprised of a (latent) main process from which two branches diverge and recombine, was fitted to the simulated data, with hyperparameters optimised using type II ML estimates. Example fits are shown in Figure 1(c), with the log mean sum square error shown in used to generate data, corresponding to a HGP, *f*_0_(*t*)~*GP*(0, *K*_0_(*t,t′*), *f*_1_(*t*)~*GP*(*f*_0_ (*t*), *K*_1_ (*t*,*t′*)), *f*_2_(*t*)*~GP*(*f*_0_(*t*),*K*_2_(*t,t′*)), with squared-exponential covariance functions used throughout. A B-RGP was fitted to the data with hyperparameters initialised as *t_b_1__* *=* −4, *t_r_1__*= 4, log s*_b_1_, b_2_r_1_ r_2__* = 0.5, and log *σ_n_*= 0.2, and all other hyperparameters initiated as *θ*_*i*_~ *U*(0.1,1). In this case, there exists a model mismatch between the data, which has no explicit branching or recombination, and the branch-recombinant covariance function used for inference. Nevertheless, we note that informally, if the branch point occurs much earlier than the first data point and the recombinant point occurs much later than the last data point, the behaviour over the range of observations is identical to that of a HGP, *f*_1_(*t*)~*GP* (0,*K*_*b*_1__(*t,t′*) + *K*_*b*_2__(*t,t′*)), *f*_2_(*t*)*~GP* (0,*K*_*b*_1__(*t,t′*) + *K*_*b*_3__(*t,t′*)). Tuning of the branch/recombination time hyperparameters should therefore allow a good fit over the regions of observation despite the model mismatch. In Supplementary Figure 3(a) we plot example fits to the function using a B-RGP, BGPs, IGPs, a JGP and splines. In Supplementary Figure 3(b,c) we indicate the sum of squared errors and negative log marginal likelihoods. As expected, B-RGPs and IGPs were more accurate than other approaches, due to the increased flexibility to fit the two processes, rather than fitting the general underlying trend. In most cases B-RGPs performed comparably to IGPs, although in a few instances the B-RGP Figure 1(d). Again, branch-recombinant GPs outperformed all other methods, with branching GPs DEtime performing next best.

To test for robustness to model mismatch, we used B-RGPs on two other datasets. In dataset 3, a non-branching, three-component process was appeared to suffer from numerical instability and failed to converge, with the resulting mean SSE and negative log marginal likelihood distributions heavy tailed and not as favourable as for IGPs. These results suggest that B-RGPs offer comparable performance to IGPs, although performance depends on sensible initialisation of hyperparameters.

To further evaluate the effect of model mismatch, we fit to data from a single, noisy process. Specifically, we used the same three-component HGP as in dataset 3, with noisy observation data generated from the first process only i.e., representing two replicates 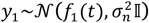, 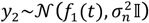. As before, we fitted the data using a three-component branch-recombinant process, with squared-exponential co-variance function assumed for all branches, and hyperparameters initiat-ed *t*_*b*_1__=−4, *t*_*r*_1__ = 4. Informally, we note that, despite the model mismatch, when branching and recombination both occur much earlier than the first observation, or much later than the last observation, the fit over the range of observations should correspond to that of a JGP with covariance function corresponding to that of the main branch process, *f*_1,2_(*t*)0~*GP*(0,, *K*_0_(*t,t′*)). In Supplementary Figure 4(a) we indicate example fits to the function using a B-RGP, IGPs and a JGP, whilst in Supplementary Figure 4(b, c) we indicate the SSE and negative log marginal likelihood. In general, both the B-RGP and JGP outperform the other approaches.

Finally, we performed inference on a four-branch system, in which we have one latent basal branch, from which two intermediate latent branches emerge. For comparison, we evaluate the sum squared error for the B-RGP, IGPs a JGP, and splines, with the results indicating B-RGPs provide better overall performance (Supplementary Figure 4).

Together, analysis of datasets 1 - 4 indicate B-RGPs offer superior performance compared to other approaches when the underlying data is branch-recombinant, with good ability to estimate the timing of bifurcations. Crucially, all hyperparameters can be optimised directly using type II ML. Therefore, branch and recombination time hyperparameters can be tuned, which, due to their limiting behaviour, means that they can gracefully cope with datasets where no branching structure exists, provided hyperparameters are sensibly initialised.

### 3.1. Inference for partially labelled datasets

In our previous examples, inference relied on the existence of explicit branch labels. In some cases, however, branch labels may be incomplete or missing entirely. For example, in a collection of single cell transcriptomics data there may be various cell types, including some that cannot be unambiguously assigned to a particular branch *a priori.* We can attempt to infer the branch labels, **z**, using Markov chain Monte Carlo (MCMC). Here we assume partially labelled data, with a subset of branch labels know, and the remainder unknown, denoted **z** = [**z**^(labelled)^**,z**^(unlabelled)^]. When branch labels are known, they can be fixed, whilst unknown branch labels are initialised stochastically, and updated via a Gibbs sampler, similar to the usage in Stegle et al. (2010). For an n-component branching process, the unknown label for cell *i*, is Gibbs sampled conditional on the observation data and branch assignment of all other cells:

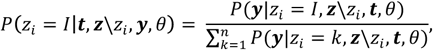

with hyperparameters updated conditional on all branch labels using hybrid Monte Carlo (HMC):

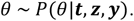

To test the accuracy of our B-RGPs on partially labelled data we generated observations from the simple branching process outlined in Sup-plementary Section 2. We first generated a set of test input locations, *t*~*𝒩*(0,5𝕀), with observation data generated as noisy instances of the process. We then attempted to infer branch labels and hyperparameters within an MCMC scheme, with labels updated via Gibbs sampling, and hyperparameters sampled using Hybrid Monte Carlo. A subset of data points, *n*, were assigned the correct branch label, where *n/N* G [0,0.1,0.25,0.5] and *N* indicates the total number of observations, with the remaining data points randomly assigned to either branch with equal probability and updated within the MCMC. Five randomly initiated runs were used, with 20,000 steps in the MCMC chain, and the first 5,000 discarded for burn-in. An example of the initial branch assignment is shown in Supplementary Figure 6(a), with red indicating data points initially assigned to branch 1, and blue assigned to branch 2. An example fit (and updated branch labels) is shown for step 20,000 in Supplementary Figure 6(b). The accuracy of classification is summarised using receiver operating characteristic (ROC) curves in Supplementary Figure 6(c, d). We note good overall ability to infer branch labels even for the unlabeled case.

## 4 Arabidopsis thaliana transcriptional branching in response to Pseudomonas syringae

To evaluate the utility of B-RGPs on a genome scale applications, we used our framework to investigate transcriptional branching in model plant organism *Arabidopsis thaliana* in response to infection with hemibiotrophic bacterial pathogen *Pseudomonas syringae.* Recent studies by Lewis *et al.*, 2015 (GEO GSE56094) have provided highly temporally resolved transcriptional datasets for Arabidopsis following inoculation with disease-causing *Pseudomonas syringae* pv. tomato DC3000, and a disarmed mutant strain hrpA using bulk microarray measurements. Whilst both strains efficiently infect the plant cells, the DC3000 variant also delivers 28 effector proteins that subvert the plant’s immune response; the disarmed hrpA mutant lacks the apparatus for effector delivery and thus elicits a classical immune response. Yang *et al.* (2016) developed a two-component branching GP to investigate transcriptional bifurcations between time series of hrpA- and DC3000-inoculated cells. Here we extend this analysis by simultaneously deciphering the branching structure that exists between all 3 time series (mock/control, virulent (DC3000) and innate immune (hrpA) responses).

For each gene in the three datasets, we consider a number of possible branching structures: hrpA branches from the control, with DC3000 branching from hrpA at a later point (Group 1), or hrpA and DC3000 independently branch from the control (Group 2), which collectively represent immune response genes that are targeted by effectors; DC3000, but not hrpA, branches from control (Group 3), representing host susceptibility genes that have been targeted by effectors; hrpA, but not DC3000, branches from the control (Group 4), likely representing immune genes that have been targeted by effectors prior to their natural immune response times; both DC3000 and hrpA jointly branch from the control, but not from one another (Group 5), representing core immune response genes not targeted by effectors; no branching exists (Group 6), representing genes unaffected by plant immunity or pathogen virulence strategies. Example expression patterns of individual genes from each of the six groups are shown in Figure 2(a).

For Group 1, we assume that the hrpA-infected time series branches from the mock-infected time series, with the DC3000-infected time series branching from hrpA-infected:

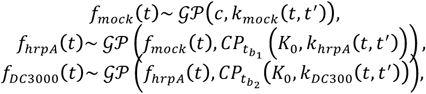

where observation data was assumed to be a noisy instances of these functions e.g., *y*_*mock*_(*t*) *=f*_*mock*_(*t*) +*ε.* For Group 2, we have hrpA-infected and the DC3000-infected time series independently branching from the mock-infected time series:

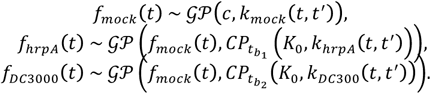

Collectively Groups 1 and 2 should represent immune response genes targeted by effectors, and therefore associated with the onset of disease. For Group 3, we have mock-infected and hrpA-infected datasets drawn from an identical process, with the DC3000-infected branching from this:

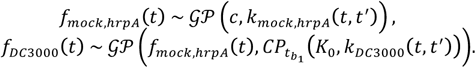

This group represents genes not associated with the immune response that are nonetheless targeted by effectors, and may therefore represent those functioning in metabolism. For Group 4, we have mock-infected and DC3000-infected datasets drawn from an identical process, with the hrpA-infected branching from this:

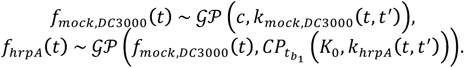

These genes likely reflect downstream immune response genes that are targeted very early by effectors, before their usual time of immune activation. For Group 5, we have hrpA-infected and DC3000-infected datasets drawn from an identical process that branches from mock-infected:

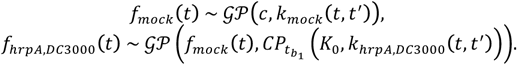

These genes represent immune response genes not targeted by effectors. Finally, for Group 6, we have all datasets drawn from an identical process:

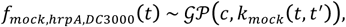

representing genes that are unbranched i.e., not differentially expressed. Because these datasets correspond to bulk observations from microarrays with well-defined measurement times we assumed smooth functions throughout, and therefore, in all cases, the covariance functions were taken to be squared-exponentials e.g., 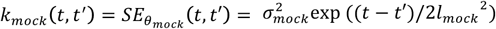 where 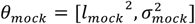 denotes a set of mock dataset-specific hyperparameters, and hyperparameters were optimised to their ML or MAP values. We assumed the following prior distributions: the first branch time was Gamma distributed, *t*_*b*_1__~Γ(2,2), with the second branch also Gamma distributed, *t*_*b*_1__~Γ(4,2), and the change-point transition rate was Gaussian distributed, *s* ~ 𝒩(0,0.5). All other hyperparameters were optimised to their ML values. Finally, we selected the optimal group based on the Bayesian information criterion (BIC).

In Supplementary Figure 7(a), we indicate the branch time between control and hrpA time series using B-RGPs versus that obtained using the Gaussian process two-sample (GP2S; Stegle et al. (2010); Supplementary Figure 7(b)). Here the GP2S approach incorrectly identified a peak perturbation time at *t* = 0, before Arabidopsis could mount an immune response. This peak was notably absent in our B-RGP approach. To further gauge the accuracy of our approach, we compared the estimated branch times between hrpA and DC3000 using B-RGPs (Supplementary Figure 6(c)) to that obtained using the perturbation times previously estimated in Yang et al. (2016) (Supplementary Figure 6(d)). The analysis in Yang et al. (2016) provide 90% confidence intervals for branch time estimates, and we note that our MAP estimation falls within these bounds in 67% of cases. Of the remaining genes, 27% of our MAP estimates lie to the right of the confidence bounds, and 5% to the left of the confidence bounds, suggesting that our approach has a tendency to estimate later branch times than that of Yang et al. (2016). This is likely due to differences in the prior distributions over branch times. Indeed, if we plot the estimated branch time using our method versus the PT approach for the 27% of genes noted above, we see a strong correlation (R = 0.8507). Together, these results suggest that, although there is good agreement between the methods for a large fraction of cases, inference of branch time in a subset of the observations may be unidentifiable.

Our results indicate that approximately 50% of genes were unperturbed by either the hrpA or DC3000 strains, in agreement with previous studies based on pairwise comparison of the time series using mixtures of Gaussian Processes (Lewis et al. 2015), where 52% of genes were identified as being differentially expressed in control versus hrpA or control versus DC3000.

Since DC3000 is known to subvert the basal immune response of Arabidopsis, we hypothesised that the expression of a subset of genes in the DC3000-infected dataset might converge fully back to control levels following delivery of effectors. To identify such genes, we also fitted a two-component branch-recombinant GP using the control and DC300-infected time series only, again using the BIC to distinguish between genes undergoing branching and recombination from those undergoing branching alone.

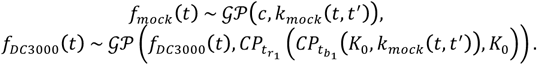

An example branch-recombinant expression profile is shown in Figure 2(b). We note that relatively few genes were identified as having their expression levels fully converge back to control levels. Of those that did, none were identified as being targets of effectors or previously implicated in the response to *P. syringae*, suggesting that the full suppression of early immune response genes to control levels is not required for infection to advance.

Gene Ontology analysis identified several highly enriched terms across the first five groups (see Supplementary Table 1), suggesting distinct biological functions relating to pathogen response and metabolic repro remodelling in this dataset, B-RGPs provided much better temporal resolution. As the effector-driven virulence programme proceeds, but prior to bacterial multiplication (5-8 hpi), there is a strong enrichment of gramming. Prior to 3 hours, the ontologies represent some of the earliest transcriptional processes targeted by effectors. Consequently, there is a diverse array of GOs represented. Notable are the combination of proteolytic, ribosome, vitamin and amino acid metabolic and transport processes. This is indicative of assembly of the processing machinery to enable effector mediated reprogramming of core cellular processes. Between 3 and 5 hours post infection (hpi) the impact of effectors was evident by the number of GOs identified, with processes associated with nuclear processes, in particular chromatin remodelling, nuclear transport, and transcription, most highly enriched. Other GO processes, such as hormone responses and primary metabolism, were, unexpectedly, less abundant. While Lewis et al. (2015) also reported evidence for chromatin terms related to adenyl ribonucleotide binding, reflecting the high energy demands at this phase of the infection process, when *Pseudomonas* effectors have suppressed immune responses and are reconfiguring the metabolism to facilitate pathogen growth.

To further investigate the nature of these groups, we looked for enrichment of known targets of effectors of various pathogens (Mukhtar et al. 2011). We first checked for enrichment of targets of *non-Pseudomonas* effectors, hypothesising that the Arabidopsis immune response to different pathogens might be conserved (Mukhtar et al. 2011). Figure 2(c, d), shows that these groups were indeed enriched. Next, we checked for enrichment of Avr and Hop effectors that are present in several strains of *Pseudomonas syringae*, and were again enriched in DC3000-responsive groups (Figure 2(c, e)).

We next looked for enrichment of genes with known pathogen-response phenotypes, using TAIR (Huala et al. 2001) to query for genes using the terms ‘Pseudomonas’, ‘Botrytis’, and ‘Peronospora’. In Supplementary Figure 8 we indicate the frequency of pathogen-response genes within various groups, and our results show a distinct enrichment for Pseudo-monas-related and Botrytis-related genes amongst the various immune responsive and disease-responsive groups.

Finally, we investigated whether inferred branch times of key regulators were predictive of branching of the direct and indirect targets of key regulators. Here we focused on WRKY11 and WRKY17, known to be amongst the earliest branching transcription factors (TFs) implicated in the *Arabidopsis* response to *P. syringae* (Journot-Catalino et al. 2006). Both genes showed branching between control and hrpA, and between hrpA and DC3000 consistent with (i) their immune-responsive expression and (ii) their suppression by DC3000 effectors. Genes that branched between control and hrpA and between hrpA and DC3000 were assigned a Euclidean distance (d) based on the position of their branch times with respect to that of WRKY 11 or WRKY 17. We then compared the distributions of these Euclidean distances for the subset of genes identified as being differentially expressed (DE) in knockout mutants of WRKY 11/17 (Journot-Catalino et al. 2006) versus the distribution of the subset of genes that were not differentially expressed (NDE) in those mutants. Our results show that DE genes had significantly smaller Euclidean distances than NDE genes (p<0.05 for WRKY 11 and p<0.005 for WRKY 17 using two-sided Student’s t-test; Figure 2(f, g)), suggesting that genes that branched at similar times to WRKY 11/17 were likely to represent a core set of genes targeted by the pathogen’s virulence strategy. WRKY 11/17 are TFs and could exert direct regulation of their targets by binding to their regulatory elements. To check this, we searched for the presence of WRKY motifs within a 1kb promoter region using FIMO (Grant, Bailey, and Noble 2011); specifically, the stringent WRKY binding site (Wbox) motif, TWGTTGACYWWWW, identified by Ciolkowski et al. (2008). Here, we looked at the frequency of Wbox motifs (p<0.0001) in sets of genes whose branch times were increasingly distal from WRKY 11. These groups were based on: (i) genes whose Euclidean distance dl, representing the closest 156 genes (see Supplementary Figure 9); (ii) genes whose Euclidean distance d<2, representing the closest 454 genes; and (iii) the closest 2000 genes. As positive and negative controls, we also included the 157 genes that were identified as DE in the WRKY 11 knockout line compared to control, and 2000 genes randomly selected from Group 6 (genes with no branching). Our results showed a clear trend of increasing frequency of Wbox motifs in sets of genes whose branch times were closest to that of WRKY 11 (Figure 2(h); see also Supplementary Table 2). Altogether, these results suggest that estimation of branch times may be useful for identifying direct and indirect targets of perturbed genes, and more generally demonstrate the efficacy of B-RGPs for extracting temporally resolved information from complex biological datasets.

## 5. Discussion

The ability to identify and quantify branching and recombination processes from systems-level measurements has a variety of important applications in the biological sciences. Here we have outlined a general framework for the composition of covariance functions that allow for the prior specification of branch-recombination processes of arbitrary complexity, both in terms of the number of branches and richness of dynamics, via simple compositional of covariance functions within a HGP framework. As well as specifying arbitrarily complex processes, all hyperparameters could be optimised via gradient based approaches, resulting in more accurate inference of branch times compared to existing approaches, although inference took slightly longer.

Here we applied B-RGPs to a time-series microarray data of *Arabidopsis thaliana* infected with a bacterial pathogen *Pseudomonas syringae.* By explicitly enumerating over all possible branch structures i.e., all 1, 2 and 3 branch structures, and using the AIC as a selection criterion, we were able to infer the branch structure for each gene. Whilst exhaustive iteration will not necessarily be possible for more complex datasets with more than three time series, we note that greedy approaches based on merging of time series could instead be used.

More generally, B-RGPs represents a flexible approach for the analysis of branching and recombination in time series datasets. This approach can be thought of as a natural extension to two-sample based approaches, allowing analysis of arbitrary numbers of time series.

Whilst here we focused on branching as a function of time, our approach is equally amenable to branching as a function of any other variable, such as expression level of a specific regulator. An intriguing possibility is therefore to incorporate B-RGPs into existing GP-based approaches for the inference of nonlinear dynamicalx systems (Penfold and Wild 2011; Penfold et al. 2012; Penfold, Millar, and Wild 2015; Penfold et al. 2015; Aijo and Lahdesmaki 2009), which would naturally allow inference of nonstationary nonlinear dynamical systems, such as temporally or spatially varying networks.

In addition, we envisage that B-RGPs could also be useful to capture transcriptional dynamics underpinning cell fate decisions from single cell transcriptomics data. For this, cells are first pseudotemporally ordered along a developmental axis using a combination of dimensionality reduction techniques and curve-fitting or graph-theoretic approaches (Trapnell et al. 2014; Bendall et al. 2014; Marco et al. 2014; Ji and Ji 2016; Setty et al. 2016). Once ordered along pseudotime, B-RGPs could capture the branching dynamics of individual genes, thus identifying the earliest molecular events controlling cell fate decisions. Alternatively, B-RGPs could be used to directly model cell fate decisions. Recent studies by (Reid and Wernisch 2016) have shown how Gaussian process latent variable modes (GPLVMs), can be used to pseudotemporally order genes along a developmental axis, with a key advantage over other pseudotime approaches: the incorporation of capture time into the inference procedure. However, due to a previous lack of treatment for branching in GP models, the approach of Reid and Wernisch (2016) did not explicitly allow for pseudotemporal ordering of datasets with branching behavior. The incorporation of B-RGPs into a GPLVM model would naturally allow for pseudotemporal ordering over branching process, whilst retaining the ability to leverage highly informative data, such as capture time.

## Acknowledgements

We thank Ufuk Günesdogan and Naoko Irie for critical reading of the manuscript, and the Surani lab for useful insights. We thank Sabine Dietmann for early discussions about branching processes. We also thank Charles Bradshaw for help with high performance computing, and the Gurdon Institute core facilities for continued support. Finally, we thank Magnus Rattray and Jing Yang for useful insights and discussion of statistical branching models.

## Supplementary materials to ‘Branch-recombinant Gaussian processes for analysis of perturbations in biological time series’

## 1 Benchmarking B-RGPs

Branching and recombinant structures can easily be encoded within a GP framework using the correct composition of covariance functions. In Supplementary Figure 1(a) we indicate example covariance functions and samples from the prior of some simple branching processes; in Supplementary 1(b) we do so for recombinant processes, and in 1(c) we do so for branch-recombinant processes.

**Supplementary Figure 1:**
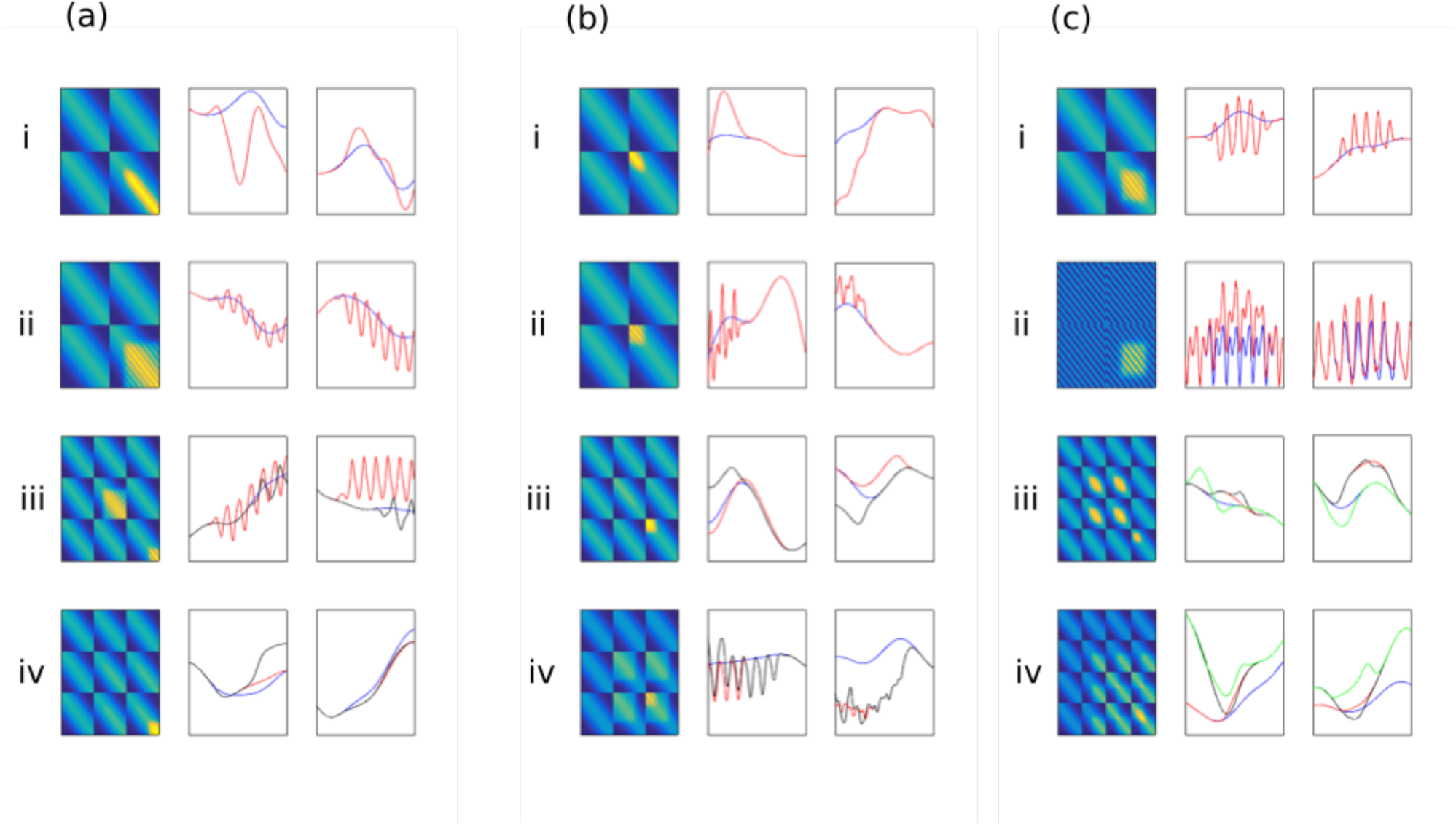
(a) GP with branching structure. Here we indicate the prior covariance matrix evaluated at uniformly incremented input times (left) and two samples from the prior distribution (middle, right). (i) A smooth function branches from a smooth base function. (ii) A periodic function branches from a smooth function. (iii) Two smooth functions (solid) branch from a smooth function (dashed). (iv) Two periodic functions (solid) branch from a smooth function (dashed). **(b) GP with recombinant structure.** (i) A smooth function converges on another smooth function. (ii) A periodic function recombines with a smooth function. (iii) Two smooth functions recombine with a smooth function. (iv) Two periodic functions recombine with a smooth function. **(c) GP with branch-recombinant structure.** (i) A periodic function branches and then recombines with a smooth function. (ii) Two periodic functions branch and then recombine with one another. (iii) A set of four smooth functions branch and recombine at various time points. (iv) A set of two functions (black and green) branch and recombine with one another, whilst a third function (red) recombines with the first two functions, and a fourth function (blue) branches from the third function and recombines with the first two

**Supplementary Figure 2:**
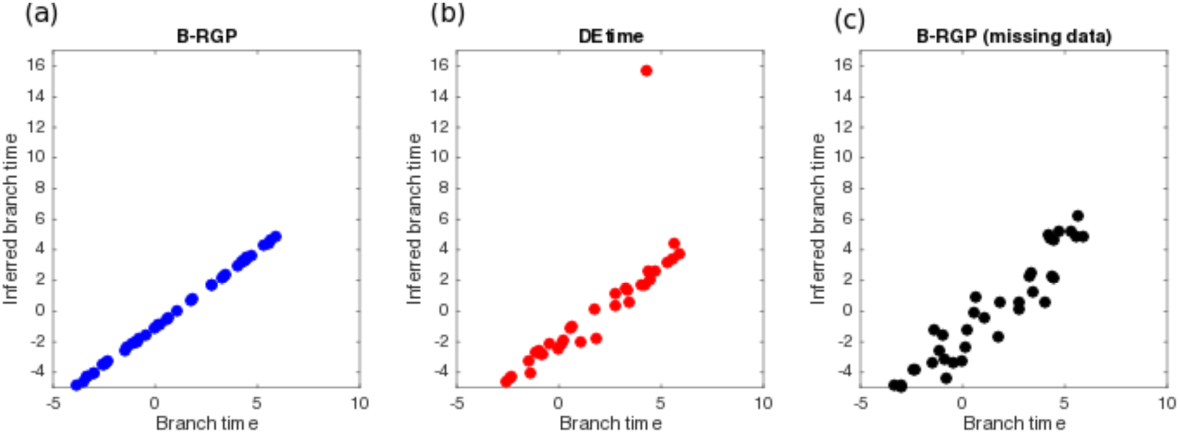
Branch time versus inferred branch time versus using a B-RGP **(a)** and the DEtime approach **(b)** for 50 randomly generated branch processes show high correlation of *R* = 0.9999 and *R* = 0.9007 respectively, indicating good ability to infer the time of bifurcations in time series. **(c)** Branch times versus inferred branch time for B-RGPs with missing data centered around the branch point (*R* = 0.9649).

**Supplementary Figure 3:**
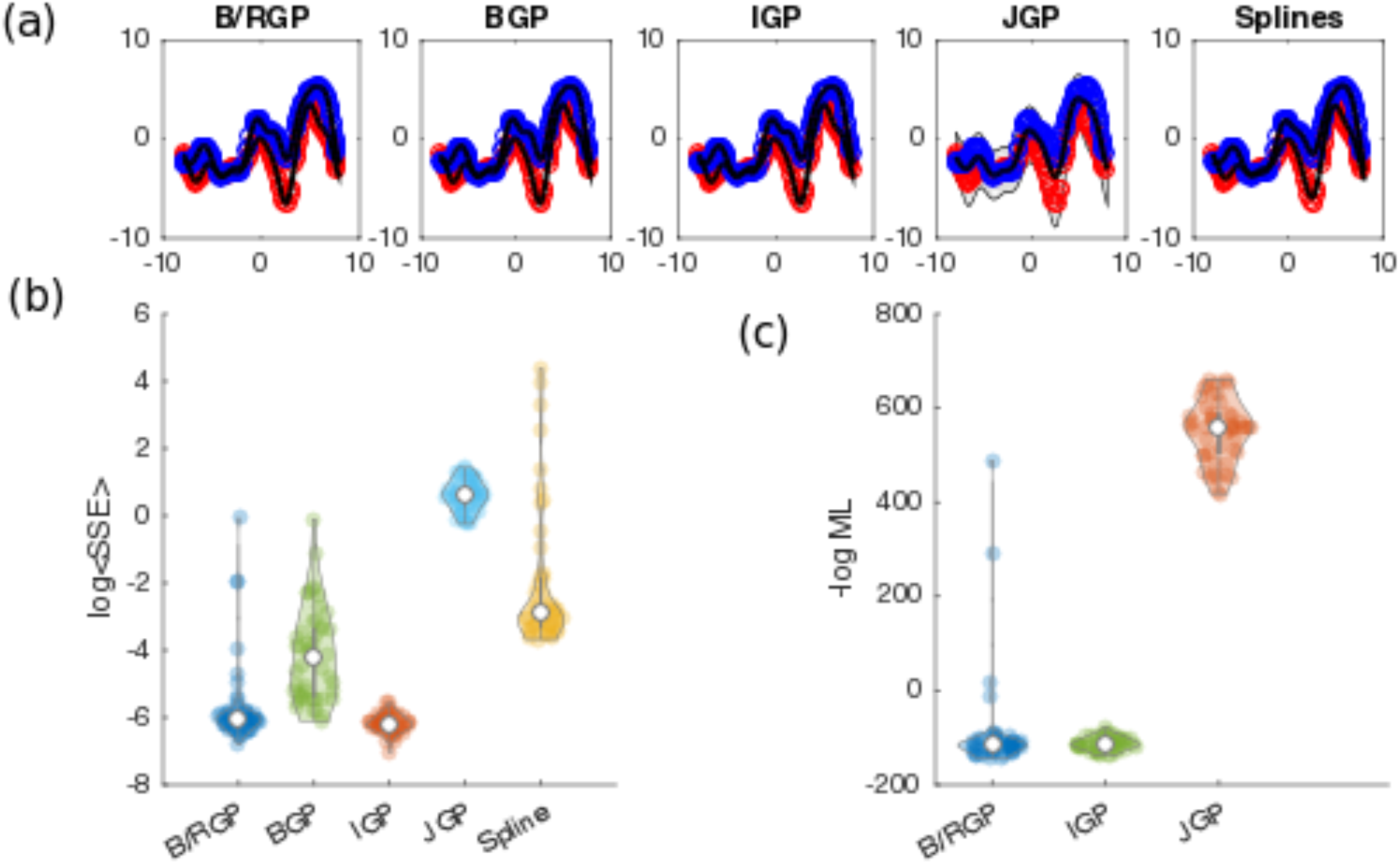
(a): Example posterior GPs fitted to a non-branching hierarchical GP using a B-R GP (left), independent GPs (middle) and a joint GP (right). **(b):** distribution of mean sum square error (left) and negative log marginal likelihood (right) for B-RGP, limiting case of a B-RGP (*), IGP and JGP. **(c) Example posterior GPs fitted to a non-branching process** (datasets represent replicated samples from a GP) using a three-component branch-recombinant GP (left), independent GPs (middle) and a joint GP (right). **(d)** distribution of mean sum square error (left) and negative log marginal likelihood (right) for B-RGP, limiting case of a B-RGP (*), IGP and JGP.

**Supplementary Figure 4:**
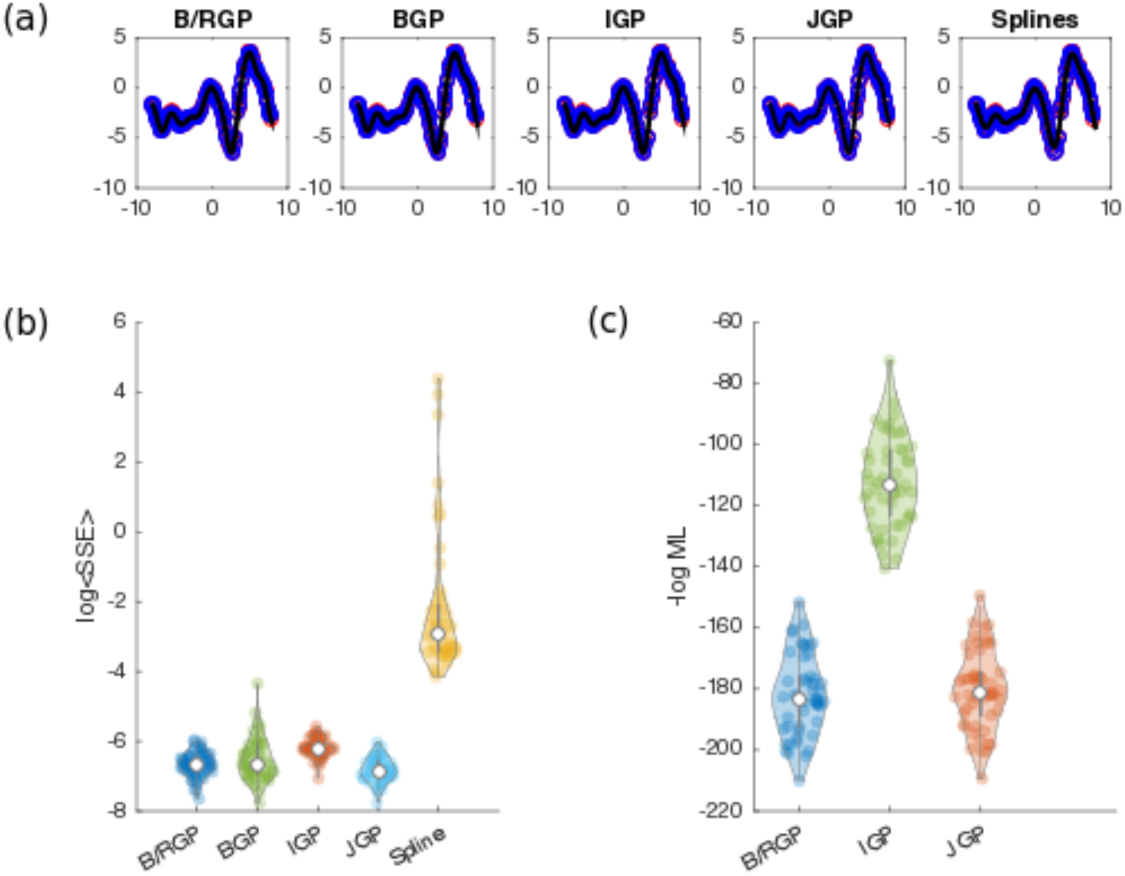
(a): Example posterior GPs fitted to a non-branching hierarchical GP using a B-R GP (left), independent GPs (middle) and a joint GP (right). **(b):** distribution of mean sum square error (left) and negative log marginal likelihood (right) for B-RGP, limiting case of a B-RGP (*), IGP and JGP. **(c) Example posterior GPs fitted to a non-branching process** (datasets represent replicated samples from a GP) using a three-component branch-recombinant GP (left), independent GPs (middle) and a joint GP (right). **(d)** distribution of mean sum square error (left) and negative log marginal likelihood (right) for B-RGP, limiting case of a B-RGP (*), IGP and JGP.

**Supplementary Figure 5:**
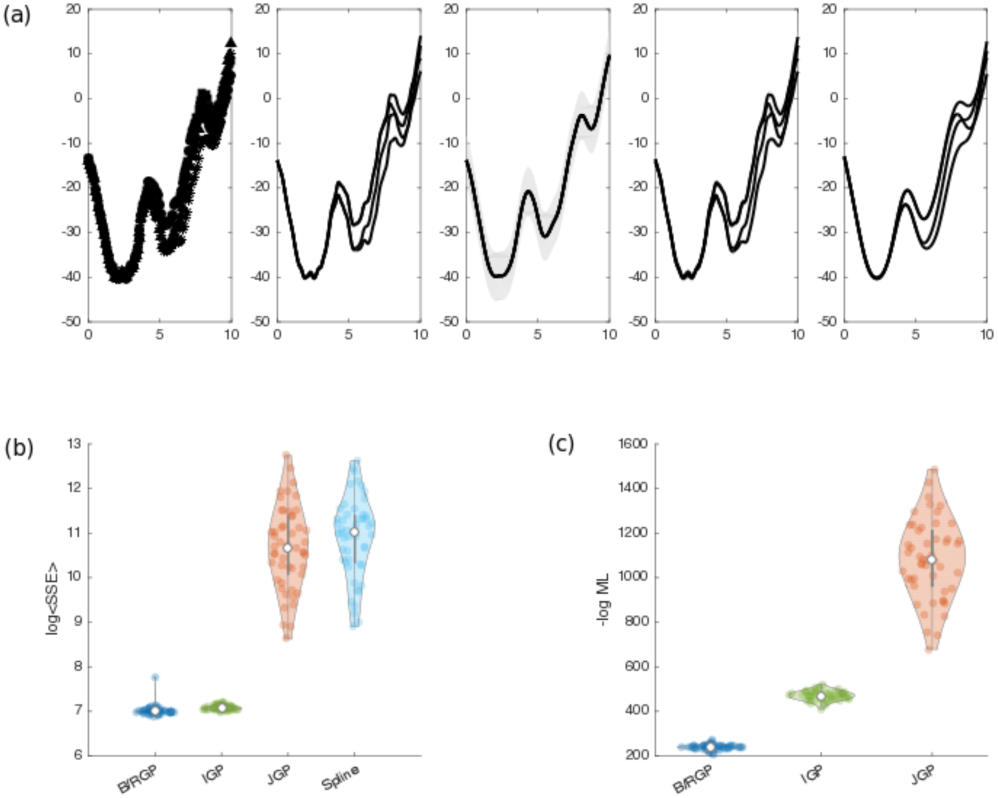
(a): Example posterior GPs fitted to a complex branching process using B-RGPs, JGPs and IGPs. Here we indicate accuracy of the inference using: (b) The log sum squared error; and (c) the log marginal likelihood. In both cases B-RGPs were more accurate than IGPs and JGPs.

## 2 B-RGPS FOR PARTIALLY LABELLED DATA

In Supplementary Figure 5 we show the accuracy of inferred B-RGPs for branching data for partially labelled datasets.

**Supplementary Figure 6:**
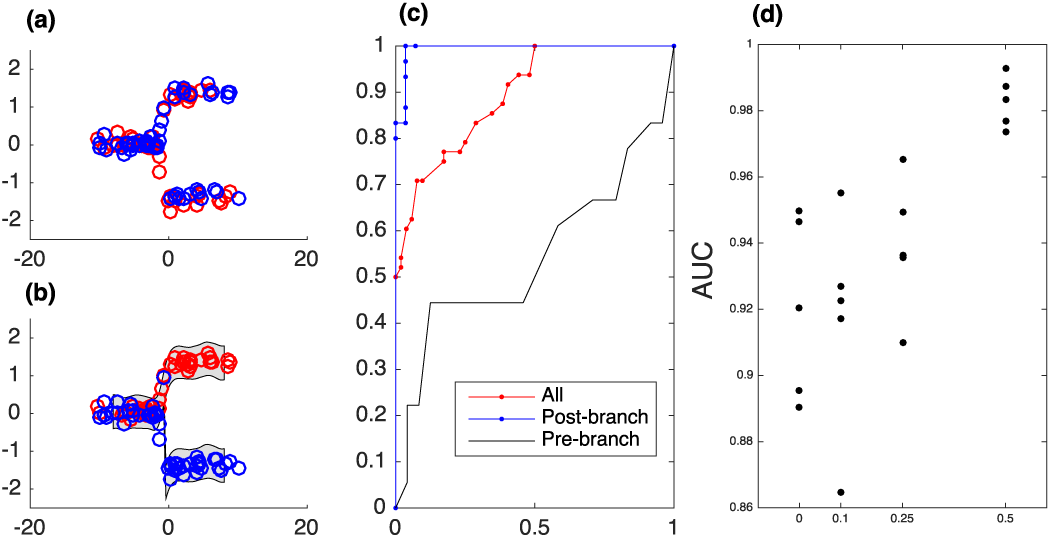
Inferring BGPs with unlabelled data. **(a)** The initial assignment of branch labels is made with a fraction initialised to the correct values, and the remainder initialised stochastically. **(b)** Unknown labels are updated via Gibbs sampling, with hyperparameters sampled every 100 steps using HMC. Here we indicate the BGP fit after 20,000 iterations. **(c)** Receiver operating characteristic (ROC) curve indicating the accuracy of inferred branch labels. **(d)** Area under the ROC curve (AUC) plotted as a function of the fraction of correctly initialised branch labels.

## 3 Comparison of B-RGPs versus other approaches for the Pseudomonas data

**Supplementary Figure 7:**
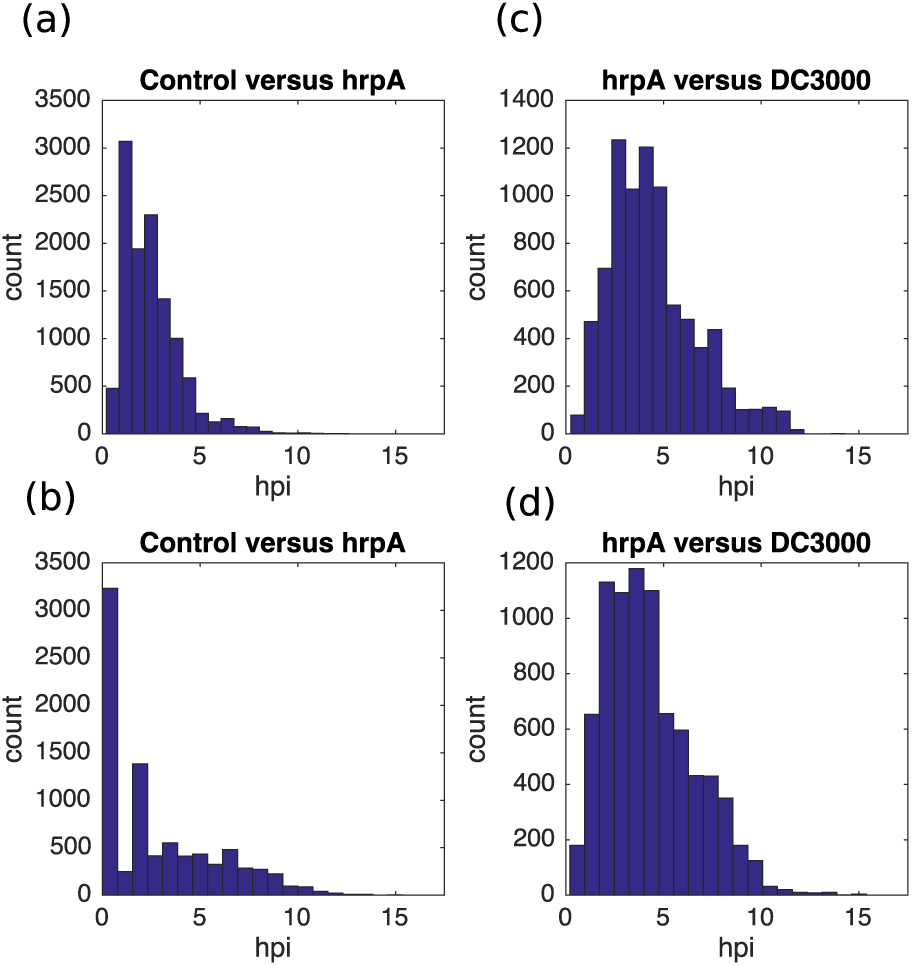
Histogram of the inferred branch times using B-RGPS. Branch times between mock-infected and hrpA-infected plants are shown for B-RGPs **(a)** and a mixture of Gaussian processes (GP2S) **(b)**. Branch times between hrpA-infected and DC3000-infeced Arabidopsis using B-RGPs **(c)** and Perturbation Time (PT) analysis **(d)**.

**Supplementary Figure 8:**
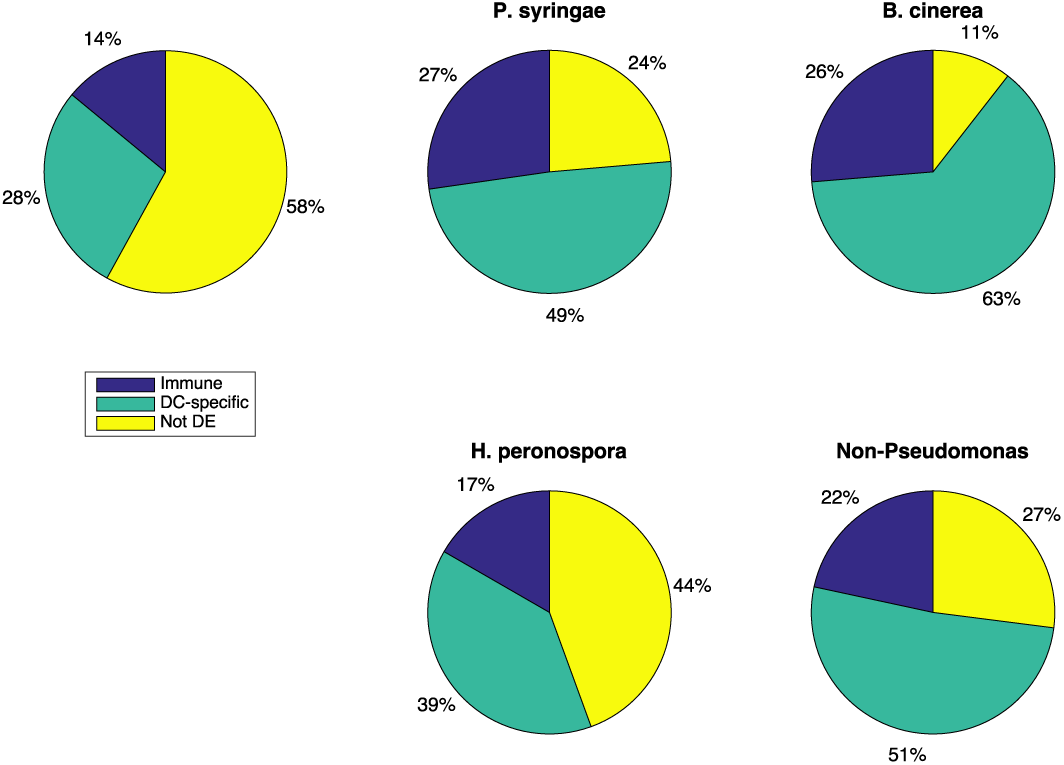
Frequency of known ‘Pseudomonas’, ‘Botrytis’, and ‘Peronospora’ genes within various groupings. Genes were first grouped according into immune-responsive (Groups 1, 2, 4 and 5), DC-specific (Groups 1, 2,3, and 4) versus not DE groups. Statistical significance was determined using a Chi-squared test on the analysis of variance between Poisson GLM fits with and without interaction terms. Here we note that immune responsive and DC-specific subgroups were both enriched for *Pseudomonas* and *Botrytis* related genes, although no enrichment for *H. peronospora* was noted.

**Supplementary Figure 9:**
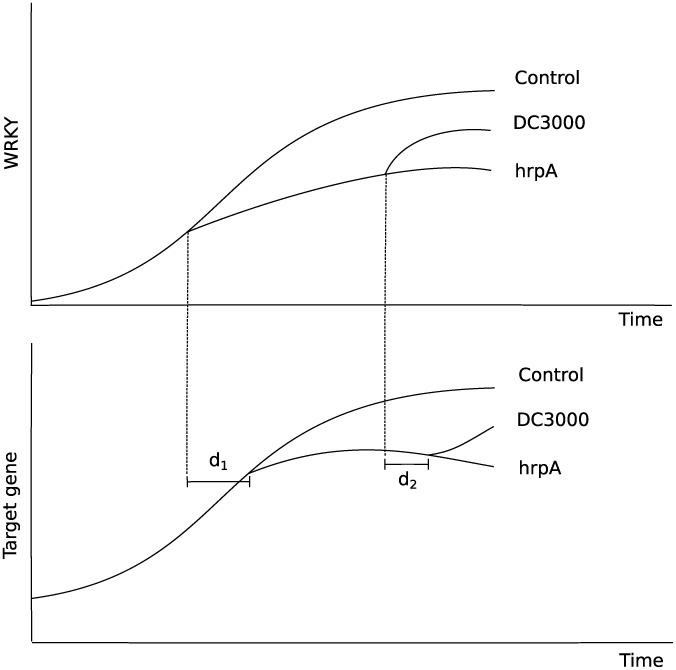
Metric used to evaluate distance of branching time for target gene to WRKY. Here we calculate the Euclidean distance between the branch times between WRKY 11/17 and all other genes in Groups 1 and 2 as *d* = |*d*_1_ + *d*_2_|.

**Supplementary Tables 1 and 2**: Available as separate spreadsheets

## References

Aijo, T., and H. Lahdesmaki. 2009. ‘Learning gene regulatory networks from gene expression measurements using non-parametric molecular kinetics’, Bioinformatics, 25: 2937–44.

Bendall, S. C., K. L. Davis, A. D. Amir el, M. D. Tadmor, E. F. Simonds, T. J. Chen, D. K. Shenfeld, G. P. Nolan, and D. Pe’er. 2014. ‘Single-cell trajectory detection uncovers progression and regulatory coordination in human B cell development’, Cell, 157: 714–25.

Boller, Thomas, and Sheng Yang He. 2009. ‘Innate Immunity in Plants: An Arms Race Between Pattern Recognition Receptors in Plants and Effectors in Microbial Pathogens’, Science, 324: 742–44.

Breeze, E., E. Harrison, S. McHattie, L. Hughes, R. Hickman, C. Hill, S. Kiddle, Y. S. Kim, C. A. Penfold, D. Jenkins, C. Zhang, K. Morris, C. Jenner, S. Jackson, B. Thomas, A. Tabrett, R. Legaie, J. D. Moore, D. L. Wild, S. Ott, D. Rand, J. Beynon, K. Denby, A. Mead, and V. Buchanan-Wollaston. 2011. ‘High-resolution temporal profiling of transcripts during Arabidopsis leaf senescence reveals a distinct chronology of processes and regulation’, Plant Cell, 23: 873–94.

Ciolkowski, I., D. Wanke, R.P. Birkenbihl, and I.E. Somssich. 2008. ‘Studies on DNA-binding selectivity of WRKY transcription factors lend structural clues into WRKY-domain function’, Plant Molecular Biology, 68: 81–92.

Friedmann-Morvinski, D., and I. M. Verma. 2014. ‘Dedifferentiation and reprogramming: origins of cancer stem cells’, EMBO Rep, 15: 244–53.

Grant, C. E., T. L. Bailey, and W. S. Noble. 2011. ‘FIMO: scanning for occurrences of a given motif’, Bioinformatics, 27: 1017–8.

Gurdon, J. B. 1962. ‘The developmental capacity of nuclei taken from intestinal epithelium cells of feeding tadpoles’, Development, 10: 622–40.

Hensman, J., N.D. Lawrence, and M. Rattray. 2013. ‘Hierarchical Bayesian modelling of gene expression time series across irregularly sampled replicates and clusters’, BMC Bioinformatics, 14: 252.

Huala, E., AW. Dickerman, M. Garcia-Hernandez, D. Weems, L. Reiser, F. LaFond, D. Hanley, D. Kiphart, M. Zhuang, W. Huang, LA. Mueller, D. Bhattacharyya, D. Bhaya, B.W. Sobral, W. Beavis, D.W. Meinke, C.D. Town, C. Somerville, and S. Y. Rhee. 2001. ‘The arabidopsis information resource (TAIR): a comprehensive database and web-based information retrieval, analysis, and visualization system for a model plant’, Nucleic Acids Research, 29.

Ji, Z., and H. Ji. 2016. ‘TSCAN: Pseudo-time reconstruction and evaluation in single-cell RNA-seq analysis’, Nucleic Acids Res, 44: e117.

Jones, J. D., and J. L. Dangl. 2006. ‘The plant immune system’, Nature, 444: 323–9.

Journot-Catalino, N., I. E. Somssich, D. Roby, and T. Kroj. 2006. ‘The transcription factors WRKY11 and WRKY17 act as negative regulators of basal resistance in Arabidopsis thaliana’, Plant Cell, 18: 3289–302.

Kalaitzis, A.A., and N.D. Lawrence. 2011. ‘A Simple Approach to Ranking Differentially Expressed Gene Expression Time Courses through Gaussian Process Regression’, BMC Bioinformatics, 12: 180–93.

Lewis, L. A., K. Polanski, M. de Torres-Zabala, S. Jayaraman, L. Bowden, J. Moore, C. A. Penfold, D. J. Jenkins, C. Hill, L. Baxter, S. Kulasekaran, W. Truman, G. Littlejohn, J. Prusinska, A. Mead, J. Steinbrenner, R. Hickman, D. Rand, D. L. Wild, S. Ott, V. Buchanan-Wollaston, N. Smirnoff, J. Beynon, K. Denby, and M. Grant. 2015. ‘Transcriptional Dynamics Driving MAMP-Triggered Immunity and Pathogen Effector-Mediated Immunosuppression in Arabidopsis Leaves Following Infection with Pseudomonas syringae pv tomato DC3000’, Plant Cell, 27: 3038–64.

Lloyd, J.R, D. Duvenaud, R. Grosse, J.B. Tenenbaum, and Z. Ghahramani. 2014. ‘Automatic Construction and Natural-Language Description of Nonparametric Regression Models’, arXiv:1402.4304.

Marco, E., R. L. Karp, G. Guo, P. Robson, A. H. Hart, L. Trippa, and G. C. Yuan. 2014. ‘Bifurcation analysis of single-cell gene expression data reveals epigenetic landscape’, Proc Natl Acad Sci U S A, 111: E5643–50.

Mukhtar, M.S., A. Carvunis, M. Dreze, P. Epple, J. Steinbrenner, J. Moore, M. Tasan, M. Galli, T. Hao, M.T. Nishimura, S.J. Pevzner, S.E. Donovan, L. Ghamsari, B. Santhanam, V. Romero, M.M. Poulin, F. Gebreab, B.J. Gutierrez, S. Tam, D. Monachello, M. Boxem, C.J. Harbort, N. McDonald, L. Gai, H. Chen, Y. He, European Union Effectoromics Consortium, J. Vandenhaute, F.P. Roth, D.E. Hill, J.R. Ecker, M. Vidal, J. Beynon, P. Braun, and J.L. Dangl. 2011. ‘Independently Evolved Virulence Effectors Converge onto Hubs in a Plant Immune System Network’, Science, 333: 596–601.

Penfold, C. A., V. Buchanan-Wollaston, K. J. Denby, and D. L. Wild. 2012. ‘Nonparametric Bayesian inference for perturbed and orthologous gene regulatory networks’, Bioinformatics, 28: i233–41.

Penfold, C. A., J. B. Millar, and D. L. Wild. 2015. ‘Inferring orthologous gene regulatory networks using interspecies data fusion’, Bioinformatics, 31: i97–105.

Penfold, C. A., A. Shifaz, P. E. Brown, A. Nicholson, and D. L. Wild. 2015. ‘CSI: a nonparametric Bayesian approach to network inference from multiple perturbed time series gene expression data’, Stat Appl Genet Mol Biol, 14: 307–10.

Penfold, C. A., and D. L. Wild. 2011. ‘How to infer gene networks from expression profiles, revisited’, Interface Focus, 1: 857–70.

Poincaré, H. 1885. ‘Sur l’équilibre d’une masse fluide animée d’un mouvement de rotation’, Acta mathematica, 7: 259–380.

Rasmussen, C. E., and C. K. Williams. 2006. Gaussian processes for machine learning (MIT Press).

Reid, J.E., and L. Wernisch. 2016. ‘Pseudotime estimation: deconfounding single cell time series’, Bioinformatics, 32: 2973–80.

Setty, M., M. D. Tadmor, S. Reich-Zeliger, O. Angel, T. M. Salame, P. Kathail, K. Choi, S. Bendall, N. Friedman, and D. Pe’er. 2016. ‘Wishbone identifies bifurcating developmental trajectories from single-cell data’, Nat Biotechnol, 34: 637–45.

Stegle, O., K.J. Denby, E.J. Cooke, D.L. Wild, Z. Ghahramani, and K.M. Borgwardt. 2010. ‘A Robust Bayesian Two-Sample Test for Detecting Intervals of Differential Gene Expression in Microarray Time Series’, JOURNAL OF COMPUTATIONAL BIOLOGY, 17: 355–67.

Takahashi, K., and S. Yamanaka. 2006. ‘Induction of pluripotent stem cells from mouse embryonic and adult fibroblast cultures by defined factors’, Cell, 126: 663–76.

Trapnell, C., D. Cacchiarelli, J. Grimsby, P. Pokharel, S. Li, M. Morse, N. J. Lennon, K. J. Livak, T. S. Mikkelsen, and J. L. Rinn. 2014. ‘The dynamics and regulators of cell fate decisions are revealed by pseudotemporal ordering of single cells’, Nat Biotechnol, 32: 381–86.

Windram, O., P. Madhou, S. McHattie, C. Hill, R. Hickman, E. Cooke, D. J. Jenkins, C. A. Penfold, L. Baxter, E. Breeze, S. J. Kiddle, J. Rhodes, S. Atwell, D. J. Kliebenstein, Y. S. Kim, O. Stegle, K. Borgwardt, C. Zhang, A. Tabrett, R. Legaie, J. Moore, B. Finkenstadt, D. L. Wild, A. Mead, D. Rand, J. Beynon, S. Ott, V. Buchanan-Wollaston, and K. J. Denby. 2012. ‘Arabidopsis defense against Botrytis cinerea: chronology and regulation deciphered by high-resolution temporal transcriptomic analysis’, Plant Cell, 24: 3530–57.

Yang, J., C.A. Penfold, M.R. Grant, and M. Rattray. 2016. ‘Inferring the perturbation time from biological time course data’, Bioinformatics, 32: btw329.

Zawadzka, M., L. E. Rivers, S. P. Fancy, C. Zhao, R. Tripathi, F. Jamen, K. Young, A. Goncharevich, H. Pohl, M. Rizzi, D. H. Rowitch, N. Kessaris, U. Suter, W. D. Richardson, and R. J. Franklin. 2010. ‘CNS-resident glial progenitor/stem cells produce Schwann cells as well as oligodendrocytes during repair of CNS demyelination’, Cell Stem Cell, 6: 578–90.

